# Combined multi-color immunofluorescence staining and spatial in situ mRNA expression analysis identifies potential fibrosis drivers in acute lymphoblastic leukemia

**DOI:** 10.1101/2025.08.29.673047

**Authors:** Sandro Bräunig, Carl Dencker, Dang Nghiem Vo, Rong Fan, Alba Lillo Sierras, Jens Enoksson, Anne Hultquist, Hongzhe Li, Stefan Scheding

## Abstract

Acute lymphoblastic leukemia (ALL) is the most prevalent childhood cancer. Bone marrow (BM) fibrosis in ALL has been associated with adverse outcomes, however, little is known about the mechanisms that cause fibrosis in ALL.

Therefore, we established a novel and advanced analysis method by combining multi-color immunofluorescence staining with in-situ RNA expression analysis (RNAscope^®^) investigate the spatial expression of putative fibrotic drivers in ALL bone marrows.

We analyzed standard BM biopsies from pediatric ALL patients. Sequential 5-color immunofluorescence (IF) staining with CD45, CD271, CD31, CD34 and DAPI was used to identify different BM cell types. Combined RNAscope^®^ and IF staining was established for spatial mRNA expression analysis of transforming growth factor beta 1 (*TGFB1*) and platelet-derived growth factor alpha 1 (*PDGFA1*), which are known to play major roles in primary myelofibrosis (PMF). PMF and normal BM samples served as controls.

As expected, ALL bone marrows showed high cellularities and prominent populations of blast cells. CD271^+^ MSC density was increased in ALL and was associated with fibrosis in a similar manner as observed for PMF. *TGFB1* and *PDGFA1* expression was considerably increased in ALL megakaryocytes (MKs) compared to PMF patients and normal controls. Furthermore, MK *TGFB1* and *PDGFA1* expression intensities in fibrotic ALL correlated with fibrosis grade. *TGFB1* and *PDGFA1* were also expressed in leukemic blasts, however at lower intensities compared to ALL MKs.

Taken together, advanced in-situ RNA and IF staining not only revealed increased expression of *TGFB1* and *PDGFA1* in fibrotic pediatric ALL, but also identified ALL blasts and MKs as their cellular origin at the single cell level. These novel data strongly suggest a role of these cytokines as potential fibrosis drivers in ALL. More broadly, our findings demonstrate that combined RNA and surface marker analysis is a powerful tool to provide new and valuable insights into bone marrow pathophysiology.

## Introduction

Acute lymphoblastic leukemia (ALL) is the most prevalent childhood cancer with the highest incidence occurring at very young ages. In adults, ALL is rare and the median age is approximately fifty years [1]. The efforts of cooperative multi-center groups have led to significant advancements in ALL therapy, especially for children. Generally, childhood ALL shows high survival rates, exceeding 98% in specific patient subsets [2]. Nonetheless, despite the remarkable progress in modern ALL therapy, there remains a considerable proportion of patients who do not respond optimally to treatment, resulting in an unfavorable prognosis.

Growing evidence suggests that changes in the hematopoietic microenvironment (HME) can influence treatment response. Among HME disturbances, BM fibrosis stands out as the most severe and recognizable form, characterized by excessive deposition of reticulin fibers. Accordingly, BM fibrosis has been linked to unfavorable outcomes in ALL, and high reticulin fiber density at diagnosis has been associated with elevated levels of therapy-surviving leukemia cells (minimal residual disease, MRD levels) in B-cell precursor ALL [3]. While not all studies were able to confirm the prognostic significance of fibrosis in ALL [4], it is widely agreed that BM fibrosis can be a significant and prevalent feature in ALL [5]. Recent data even shows that fibrosis in ALL is related to poor response to CAR-T cell treatments in children [6].

The mechanisms that cause fibrosis in ALL have not been investigated thus far. In contrast, more is known about the pathophysiology of primary myelofibrosis (PMF). PMF is classified as one of the myeloproliferative neoplasms (MPNs) and is considered as the prototype of a fibrotic malignancy affecting the bone marrow (BM). In addition to fibrosis, typical BM composition changes in PMF have been described in detail and include a characteristic higher presence of abnormal megakaryocytes and a reduction of erythropoiesis [7, 8], In contrast, ALL bone marrows usually show high frequencies of leukemic blast cells and reduced numbers of megakaryocytes, and, consequently, low numbers of circulating platelets [7, 9].

In PMF, driver mutations cause abnormal JAK-STAT signaling which plays a crucial role in the fibrotic process and results in the activation of NFκB (Nuclear factor kappa-light-chain-enhancer of activated B cells), the central transcriptional regulator of a wide array of inflammatory cytokines including interleukin (IL)−6, IL-1β, IL-8, tumor necrosis factor (TNF)- α, transforming growth factor beta 1 (TGFB1) and others [10, 11]. Also, elevated levels of platelet-derived growth factors (PDGF) have been linked to BM fibrosis [12]. It is worth noting that PDGFα is most prominently expressed in megakaryocytes, while PDGF receptors are predominantly found in activated fibroblasts [13-15]. Furthermore, TGF-β signaling in MSCs has been identified as an induction driver in myelofibrosis. Consequently, PDGFs and TGFBs are believed to play key roles in the proliferation of bone marrow stromal cells in fibrosis.

In this study, which is the first to investigate fibrosis mechanisms in pediatric ALL, we established combined in-situ RNA expression analysis (RNAscope^®^ ISH) with multi-color immunofluorescence (IF) staining to generate high-resolution five-color images. This enabled spatial analysis of cellular BM elements and concomitant cytokine expression at the single cell level. Using this advanced analytical approach, we investigated the expression of critical fibrosis factors in ALL patient BM biopsies to elucidate a possible involvement in fibrosis initiation and progression.

Interestingly and unexpectedly, we found that *TGFB1* and *PDGFA1* expression was notably increased in ALL megakaryocytes (MKs) and leukemic blasts whereas PMF MKs and control samples showed similar expression levels. Our data thus provide novel first insights into ALL fibrosis pathophysiology that will motivate future research aiming to develop potential anti-fibrotic strategies in ALL that may have the potential to improve response to treatment.

## Materials and Methods

### Human bone marrow biopsies

Formalin-fixed paraffin-embedded (FFPE) BM biopsies from the iliac crest bone of pediatric patients with Philadelphia negative B-ALL with fibrosis grades 0 to 2 (n=9) were analyzed. ALL marrows were compared with BM biopsies from PMF patients (n=3) and controls (n=3). Control biopsies were from lymphoma patients undergoing diagnostic BM biopsies which were diagnosed as normal by routine pathology. Patient characteristics are provided in Table 1. BM specimens were obtained from the Division of Pathology, Laboratory Medicine Skåne, Lund, Sweden. Five µm thick sections were cut from paraffin blocks and transferred to slides. Regions of interest (ROI) were defined by analyzing a HE-stained slide from each biopsy and included endosteal and perivascular areas. The sampling of patient material and sample preparation was approved by the institutional review committee in Lund (Regionala Etikprövningsnämnden i Lund, approvals no. 2014/776 and 2018/86) and all procedures were performed following the Helsinki Declaration of 1975, as revised in 2008.

### Conventional BM staining

Sections were deparaffinized in neo-clear (Merck, Germany), rehydrated in decreasing ethanol concentrations and distilled water, and stained using Hematoxylin/Eosin solution (Merck, Germany). After staining sections were dehydrated in increasing ethanol concentrations and finally neo-clear (Merck, Germany), followed by coverslip mounting. Fibrosis staining was performed using the modified Gordon and Sweet’s method [16].

### Sequential IF staining

Sequential IF staining was performed as described previously [17]. Briefly, paraffin sections were melted to improve adhesion and deparaffinized in neo-clear, rehydrated, and subjected to antigen retrieval. Specimens were stained with CD34-APC antibody (RRID: AB_2811342) followed by counterstaining with DAPI. Slides were scanned on the Olympus VS120 slide scanner. After the first scanning round, slides were bleached and stained with CD31-AF647 (RRID: AB_2801330). After a second round of scanning, the bleaching step was repeated before overnight incubation with primary CD271 (RRID: AB_2152546) and CD45 (RRID: AB_611377) antibodies. Secondary antibody staining with goat anti-mouse IgG-AF647 (RRID: AB_2338914) and goat anti-rabbit IgG-AF488 (RRID: AB_2338046) was then performed, and slides were scanned a final time for a total of three scans. Sequential scans of each slide were overlayed based on nuclei localization using the ZEISS arivis Pro software (ver. 4.0, Carl Zeiss Microscopy Software Center Rostock GmbH, Germany). For direct identification of megakaryocytes and ALL blasts, additional slides were stained with rabbit anti-human CD31 monoclonal antibody (RRID: AB_222783) and mouse anti-human CD34 monoclonal antibody (RRID: AB_306607). Secondary antibody staining was performed using goat anti-rabbit IgG AF-568 (AB_143157) and goat anti-mouse IgG-AF488 (AB_2534069). Samples were counterstained with DAPI and slides were scanned using a Leica Stellaris 5 Confocal laser-scanning inverted microscope (Leica Microsystems).

### RNA probing in combination with Immunofluorescence staining

Paraffin sections were prepared, pretreated and stained with RNAscope^®^ probes for *TGFB1* and *PDGFA*1 (Hs-TGFB1-no-XMm-C2 and Hs-PDGFA-No-XMm-C1) according to the RNAscope^®^ Multiplex Fluorescent Reagent Kit v2 Assay user manual (323100-USM/Rev Date: 12172019) with following modifications: Samples were routine biopsies processed by the pathology department. Samples were fixed in 4% formaldehyde, decalcified in MoL-DECALCIFIER (Milestone, Bergamo, Italy) and dehydrated in Logos J (Milestone, Bergamo, Italy) according to the supplier’s instructions and finally embedded by hand in Histovax (Histolab Products AB, Gothenburg Sweden). Following target retrieval, slides were baked at 60°C for 30 min. RNAscope^®^ Protease Plus treatment was performed for 30 min after which the enzyme was replaced with fresh RNAscope^®^ Protease Plus and incubated for another 10 min. In the hybridization steps for AMP 1, 2, and 3 the wash step was increased to 5 min and repeated twice. HRP-C1 signal was developed with TSA Vivid™ Fluorophore 570 diluted 1:1500, while HRP-C2 signal was developed with TSA Vivid™ Fluorophore 650 diluted 1:1500. Thereafter, slides were IF co-stained for CD45 and CD271 as follows: following antigen blocking in buffer containing 10 % goat serum (Jackson ImmunoResearch, USA) and 0.1 % sodium azide (Merck, Germany) staining was performed with primary CD271 (RRID: AB_2152546) and CD45 (RRID: AB_611377) overnight at 4 °C followed by secondary antibody staining with goat anti-mouse IgG-AF488 (RRID: AB_2338046) and goat anti-rabbit IgG-AF700 (RRID: AB_2535709) at room temperature for one hour in the dark. Finally counterstaining with DAPI and mounting was done according to the RNAscope^®^ Multiplex Fluorescent Reagent Kit v2 Assay user manual. Negative controls included probes targeting the DapB gene (accession # EF191515) of the <u>Bacillus subtilis</u> strain SMY ± TSA Vivid™ Fluorophore 570 and TSA Vivid ™ 650, respectively (supplementary Figure 1).

Scanning of the samples was performed with a Leica Stellaris 5 confocal laser-scanning inverted microscope using a 40x glycerol immersion objective. For detection of *TGFB1* and *PDGFA1* expression in CD31 stained megakaryocytes and CD34 stained ALL blasts, respectively, IF staining with anti-CD31/anti CD34 and RNA scope staining, respectively, was performed on consecutive slides. Scanning of these samples was performed using a 10x objective. The scanned images were converted to OME.TIF in QuPath (v0.5.1) and overlaid using ImageCombinerWarpy, which is included in the BIOP EPFL QuPath Warpy extension (available at https://github.com/BIOP/biop-bash-scripts.git).

### Volumetric analysis of CD271 staining

Mesenchymal stromal cell (MSC) and nuclei objects were generated using relevant pipelines from the analysis panel of Arivis Vision 4D. Pipelines are a series of operations that process images to enhance images and create objects by segmentation. Objects can then be further processed, sorted, and filtered. For our analysis, we combined the Arivis Blob Finder segmenter, which registers structures based on approximate diameter with a certain probability threshold and which can automatically split touching objects by adjusting the split sensitivity. Diameter, probability threshold, and split sensitivity were adjusted based on visual assessment of the segmented objects to match them to the expected morphology of the cells of interest in the fluorescence images. Segmented objects were further filtered by the Segment Feature Filter on channel intensities, volume, and sphericity so that the resulting objects matched the morphologically defined cell types in question. Nuclei were segmented as DAPI^+^ small round objects, while MSCs were segmented as non-spherical CD271^+^ objects, and resulting objects were filtered on low intensity in the CD45 channel. The volume of segmented objects was exported from Arivis. The volume of MSCs (CD271^+^ segmented objects with low CD45 expression) was normalized by division through the volume of nuclei in the respective region. As we analyzed individual five micrometer thick sections and not stacked consecutive slides, calculated virtual volumes reflect the area of cells rather than the total volume.

### Re-analysis of published ALL scRNA-seq data for TGFB1 and PDGFA1 expression

Single-cell RNA sequencing data from primary bone-marrow samples of a B-ALL cohort (5 patients with ETV6::RUNX1, 2 patients with Philadelphia-chromosome) across different disease stages (diagnosis -> remission -> relapse) and from healthy controls (n=4) were obtained from the GEO repository (accession number: GSE134759). For more details regarding sample isolation and sequencing library preparation please refer to the original reference [18]. For pre-processing, the full dataset included 24 samples in total that were imported directly from the separate count matrices upon data retrieval. Empty droplets were removed as were droplets containing less than 500 genes. All samples were merged and further filtering was applied to remove poor-quality sequenced cells that had: i) a mitochondrial gene content of more than 20%, ii) unique UMI counts of less than 1000 or more than 60000 reads, iii) a complexity index of less than 0.78 which was calculated as complexity index = log10(number of genes detected) / log10(number of UMI detected). Merged samples were integrated with the Harmony [19] algorithm and normalized with the SCTransform method that regressed cell-cycle scores prior to downstream clustering. UMAP was subsequently performed to obtain the integrated reduced dimensional embed across all samples using the first 40 integrated dimensions. Finally, cell-type identities were annotated with SingleR [20] using a reference dataset from the Human Primary Cell Atlas (HPCA) [21]. All pre-processing and data visualization steps were done with Seurat (V5) [22] and scCustomize [23] packages.

### Spatial *TGFB1* and *PDGFA1* mRNA expression analysis

*TGFB1* and *PDGFA1* objects were generated by Arivis analysis pipelines using the intensity threshold segmenter. Nuclei were segmented using the same pipeline as for the volumetric analysis. The surface size of segmented objects was exported from Arivis for further normalization using Microsoft Excel. For calculation of normalized expression values, the total surface of all nuclei objects in the analyzed region was divided by the total surface of *TGFB1* or *PDGFA1* objects. Calculation of the ratio of *TGFB1* and *PDGFA1* expression in MK regions vs. non-MK regions was realized by dividing the surface of *TGFB1* and *PDGFA1* objects, respectively, in MK regions by corresponding surface sizes in non-MK regions while keeping region sizes identically. Intensities of PDGFα and TGFβ expression in single cells were measured using Arivis Pro (v4.2.1). Single CD271^+^ cells, CD45^+^ cells, CD45^-^CD271^-^ cells and megakaryocytes were identifed manually and the sum intensity of PDGFα and TGFβ signals in individual cells was determined. Comparison of RNA expression levels in different cell types was performed using the Kruskal-Wallis test (Prism software version 10.4.2, Graphpad).

## Results

### Multicolor IF staining of the bone marrow architecture in acute lymphoblastic leukemia

First, we applied our established sequential multicolor IF staining [17] to analyze standard clinical FFPE BM biopsies. We obtained scans of large regions of interest (ROIs) using the Olympus VS120 slide scanner applying our five-marker panel consisting of CD34, CD45, CD271, and CD31 as well as DAPI as nuclear stain.

As shown in Figure 1, we stained BM sections from controls (normal BM, PMF) and compared them with samples from ALL patients with different degrees of fibrosis. In addition to marker panel expression, cell identity was further confirmed and refined by typical morphological features, which enabled to clearly identify MSCs, ECs, MKs, Adipocytes, and CD45^+^ hematopoietic cells (Fig. 1).

**Figure 1.**
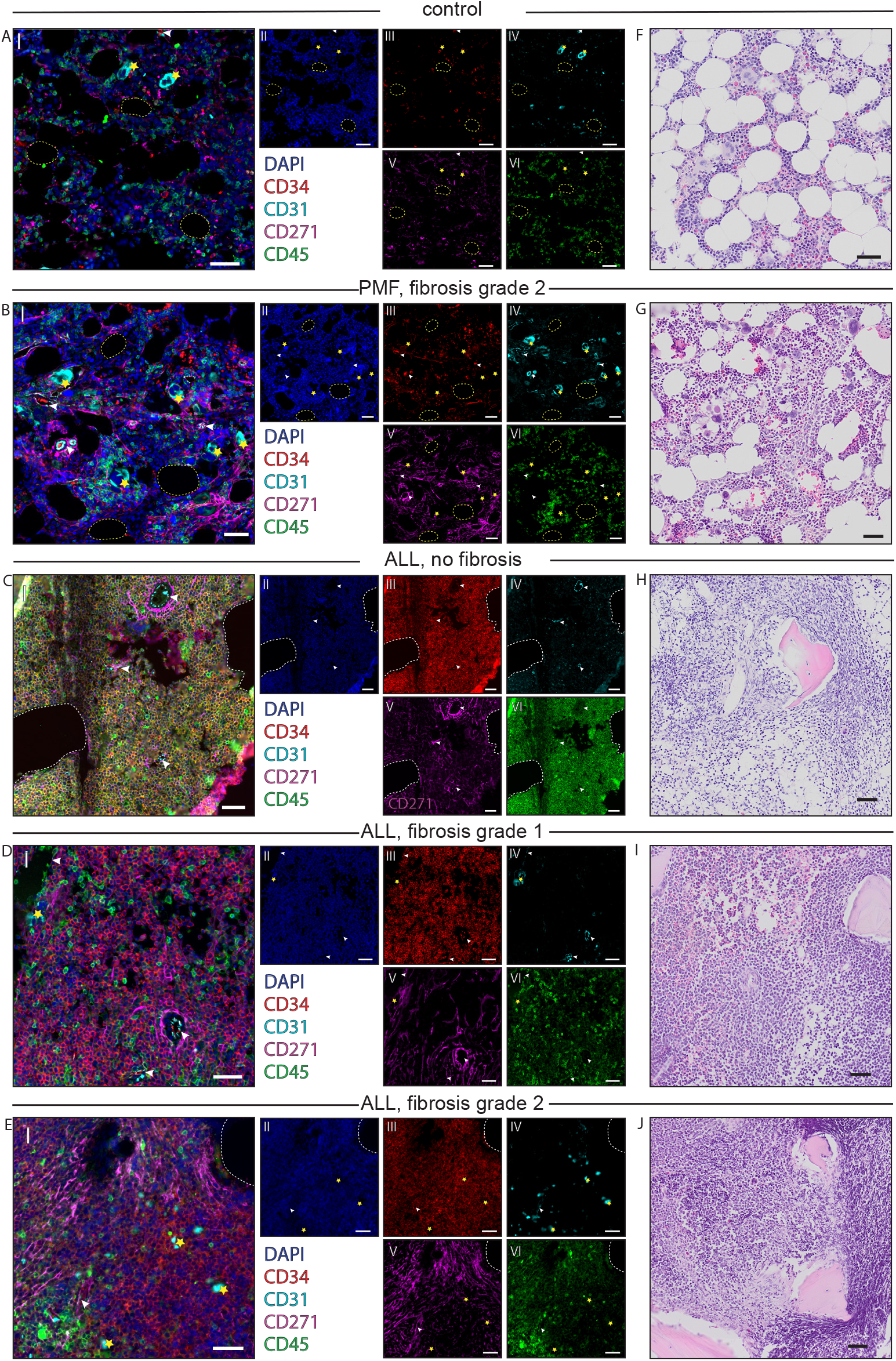
Five-color imaging of ALL, PMF, and normal control bone marrows (BMs). **(A - E)**: (I): Overlay of sequentially scanned slides from normal control BM (sample C2), PMF fibrosis grade 2 (sample F2) ALL fibrosis grade 0 (sample P2), ALL fibrosis grade 1 (sample P5), and ALL fibrosis grade 2 BM (sample P7). (II-VI): single channel images showing Nuclei stained with DAPI (blue) (II), CD34-APC in red staining ECs and HSPCs (III), CD31-AF647 in cyan staining ECs and MKs (IV), CD271 with secondary-AF647 in magenta staining for MSCs (V), and CD45 with secondary-AF488 in green staining for hematopoietic cells (VI). Yellow stars: MKs, arrowheads: ECs, white dashed lines: trabecular bone, yellow dashed lines: adipocytes. **(F-J)**: H&E staining of corresponding BM slides. Scale bars: 50 µm. For patient and sample numbers see Table 1.

Marker combinations included CD271^-^/CD31^-^/CD34^+^/45^+^ for hematopoietic stem and progenitor cells (HSPCs), CD45^-/low^/CD271^-^/CD34^low/+^/CD31^+^ for megakaryocytes (MKs), CD45^-/low^/ CD271^+^/ CD31/34^-^ for mesenchymal stromal cells (MSCs) and CD45^-^/ CD271^-^/ CD34^low/+^/ CD31^+^ for endothelial cells (ECs) [24-29].

In addition to marker expression, MSCs were thinner and elongated cells often lining other cells. ECs were identified as elongated cells lining vessels. MKs were defined as cells with a diameter of > 20µm and big lobulated nuclei. Adipocytes were identified by CD271^+^ cell membrane lining vacuoles that originally contained fat which was removed during BM biopsy processing. CD45^+^ hematopoietic cells were identified throughout the regions as smaller round cells (Fig. 1).

Trabecular bones were identified as bigger non-fluorescence areas with a low number of DAPI-positive nuclei. In several slides, bone pieces were dislocated or moved in the cutting process and washed away during staining, thus leaving empty spaces in the image (Fig. 1).

In comparison to healthy controls, MPN samples showed a tendency toward increased numbers of MKs, increased cellularity, increased MSC density especially around vessels, and fewer adipocytes (Fig. 1 A,B,F,G). The ALL samples showed an even higher cellularity and a very low number of adipocytes (Fig. 1 C-E & H-J). Furthermore, leukemic blasts (uniform small round cells) expressing surface markers CD45, CD31, and CD271 in different combinations were predominant in the ALL samples. Compared to normal controls, the density of MSCs in ALL was seemingly increased, especially around vessels, and ALL MKs were generally reduced when compared to controls and PMF (Fig. 1).

### CD271 expression correlates with fibrosis grade in ALL bone marrows

As expected, we observed increasing degrees of reticulin fibers in ALL bone marrows reflecting fibrosis grading (Fig 2, A-E). Also, we observed a clear increase in CD271^+^ MSCs in highly fibrotic ALL samples and PMF (Fig. 2B, D, E) compared to controls and non or low fibrotic ALL samples (Fig. 2A, C). Of note, CD271 expression showed a similar pattern as classic fibrosis staining, i.e. CD271 expression was found to be particularly dense close to vessels (Fig. 2B, D, E). MSC cell volumes were increased in ALL samples with fibrosis grade 2 compared to lower fibrosis ALL samples and normal controls, as were MSC volumes in PMF (Fig.2 F). Furthermore, MSC volumes were similar in high fibrosis ALL compared with PMF (Fig.2 F) but differences were not statistically significant (Kruskal Wallis test).

**Figure 2.**
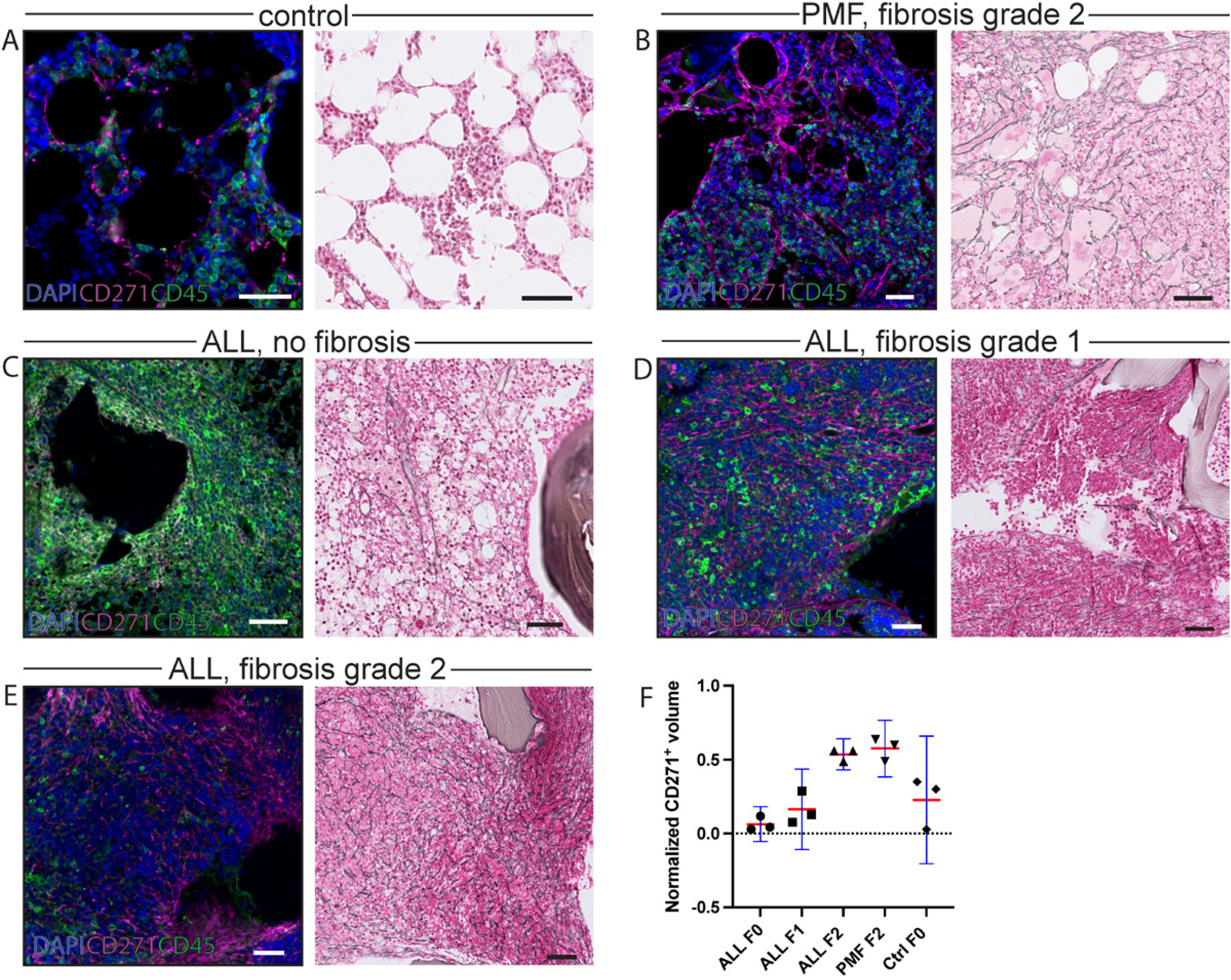
Correlation of CD271 MSC staining and fibrosis in ALL, PMF, and normal BMs. (A-E): Left: Overlay of sequentially scanned BM slides. Stainings: DAPI (blue) for nuclei, CD271 with secondary-AF647 (magenta) for MSCs, and CD45 with secondary-AF488 (green) for hematopoietic cells. Right: Fibrosis staining (Gordon & Sweet’s). (A): normal control sample C1 (B): PMF sample F1 (fibrosis grade 2) (C): ALL sample P2 (fibrosis grade 0) (D): ALL sample P6 (fibrosis grade 1) (E): ALL sample P7 (fibrosis grade 2). Scale bars indicate 50 µm. (F): Normalized total volume of CD271 single positive objects, i.e. total volume of CD271^+^ objects [μm^3^] divided by total mesh volume of DAPI^+^ nuclei [μm^3^]. Individual data (symbols), means (red horizontal lines), and 95% confidence intervals (blue) are shown. ALL F0, F1, F2 indicate ALL BMs with fibrosis grade 0, 1 and 2, respectively. PMF F2: PMF fibrosis grade 2; Ctrl F0: normal control BM, fibrosis grade 0.

### Combined IF and in situ RNA staining identified increased expression of TGFB1 and PDGFA1 in ALL blasts and megakaryocytes

The fluorescent multiplex kit RNAscope^®^ and gene-specific probes for *TGFB1* and *PDGFA1* were used for *in situ* staining of these genes as examples for important fibrosis-related mRNA targets. Combination with standard IF staining allowed for spatial expression analysis as we were able to generate scans demonstrating CD271, CD45, *TGFB1*, and *PDGFA1* expression on the single cell level in standard BM biopsy samples from patients with ALL, PMF, and hematologically normal controls (Fig. 3). The CD271/CD45/DAPI staining recapitulated the staining patterns observed with sequential IF staining as described above. *TGFB1* and *PDGFA1* expression was clearly detectable as distinctly localized bright fluorescent dots representing single mRNA molecules. In cells with higher expression, the dots fused and generated larger areas (Fig. 3). *TGFB1* and *PDGFA1* expression was comparable in PMF samples and controls (Fig. 3 A, B). In comparison, ALL samples (Fig. 3 C-E) tended to show increased overall *TGFB1* and *PDGFA1* expression (Fig 3 F, G). Visually most striking was the increased expression of *PDGFA1* and, even more clearly, expression of *TGFB1* in ALL MKs and blast cells (Fig. 3 C-E). In additional experiments, megakaryocytes were directly identified by CD31 staining, which confirmed high expression of *TGFB1* and PDGFA1 in these cells (supplementary Figure 1). Cytokine expression in ALL blasts was furthermore confirmed by co-staining with CD34 (supplementary Figure 1).

**Figure 3.**
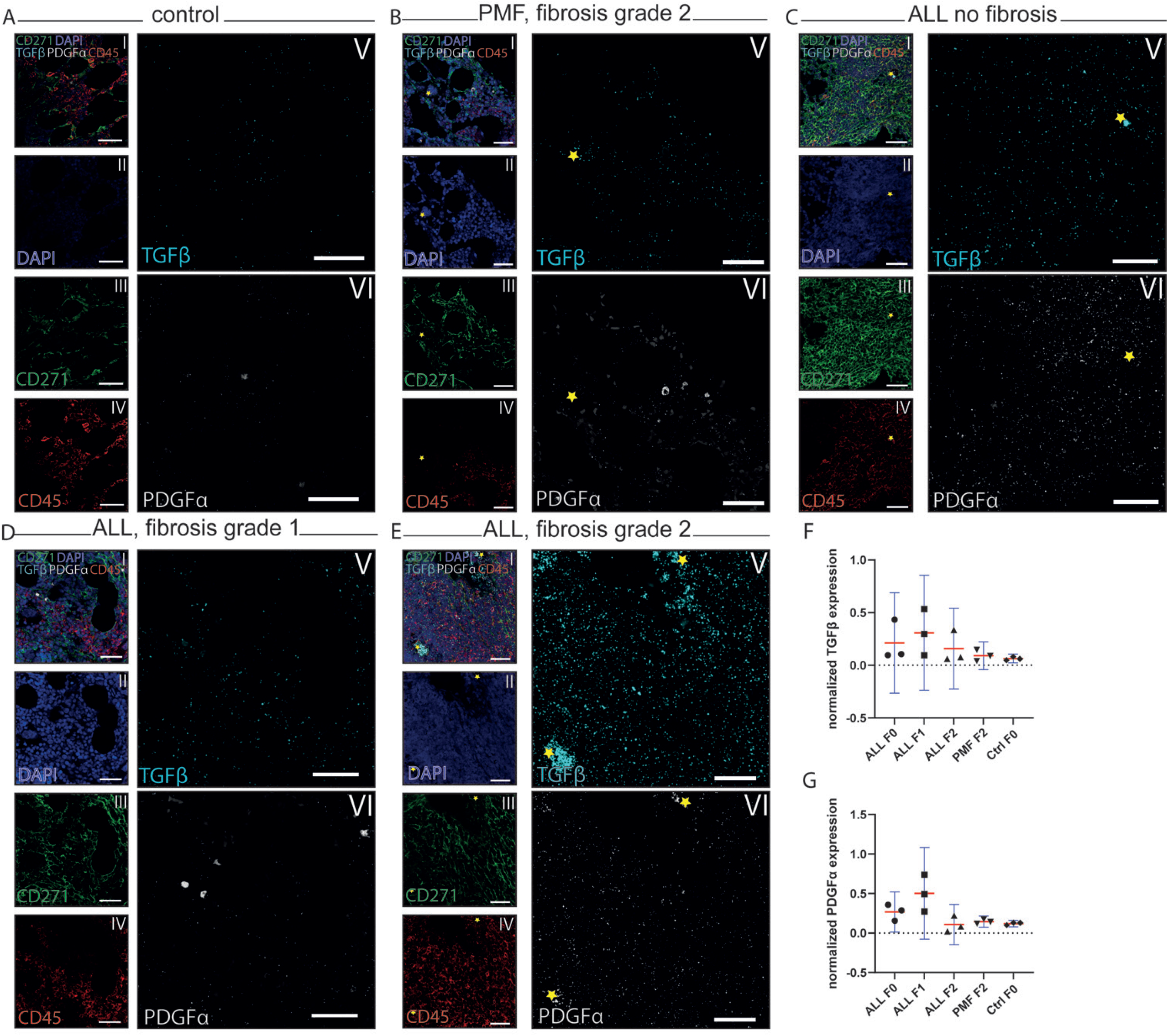
In situ expression analysis of TGFB1 and PDGFA1 in ALL, PMF, and control *BMs.* (A-E): (I): Five-channel overlay of RNAscope staining together with IF antibody staining. Yellow stars indicate megakaryocytes. (II-VI): single-channel images showing nuclei stained with DAPI (blue) (II), CD271 with secondary-AF488 in green staining of MSCs (III), CD45 with secondary-AF700 in red staining of hematopoietic cells (IV), RNAscope® probe for TGFB1-C2 coupled to TSA Vivid™ 650 in cyan (V), and RNAscope® probe for PDGFA1-C1 coupled to TSA Vivid™ 570 in gray (VI). (A): control sample C2; (B): PMF sample F2 (fibrosis degree 2); (C): ALL sample P2 (fibrosis degree 0); (D): ALL sample P5 (fibrosis degree 1); (E): ALL sample P7 (fibrosis degree 2). Scale bars indicate 50 µm. Normalized expression of TGFB1 (F) and PDGFA1 (G). Normalized expression was calculated as ratio of total DAPI ^+^ nuclei mesh areas and corresponding areas [μm ^2^] of PDGFA1 ^+^ objects and TGFB1 ^+^ objects, respectively.

### Analysis of TGFB1 and PDGFA expression in ALL-BMs using a published scRNA-seq dataset

To further investigate if *TGFB1* and *PDGFA* expression in hematopoietic cells of ALL patients could be confirmed by other methods, we reanalyzed a suitable published single-cell RNA sequencing dataset [18]. Re-annotation of the data set identified hematopoietic clusters as expected and identified differences in ALL patients at diagnosis, in remission, and at relapse, compared to healthy individuals (Fig.4 A). In diagnosis samples, a population of CD34-positive Pro-B cells emerged, representing B-ALL blasts. This population was markedly reduced in remission but emerged again in relapse samples (Fig. 4 A) as also described in the original publication [18]. Interestingly and consistent with our RNA scope data, reanalysis focusing on fibrosis-related genes clearly identified *PDGFA and TGFB1* expressing ALL blast cells at diagnosis and relapse, whereas the cells localized in the corresponding region in healthy samples showed little or no expression (Fig.4 B, C).

**Figure 4.**
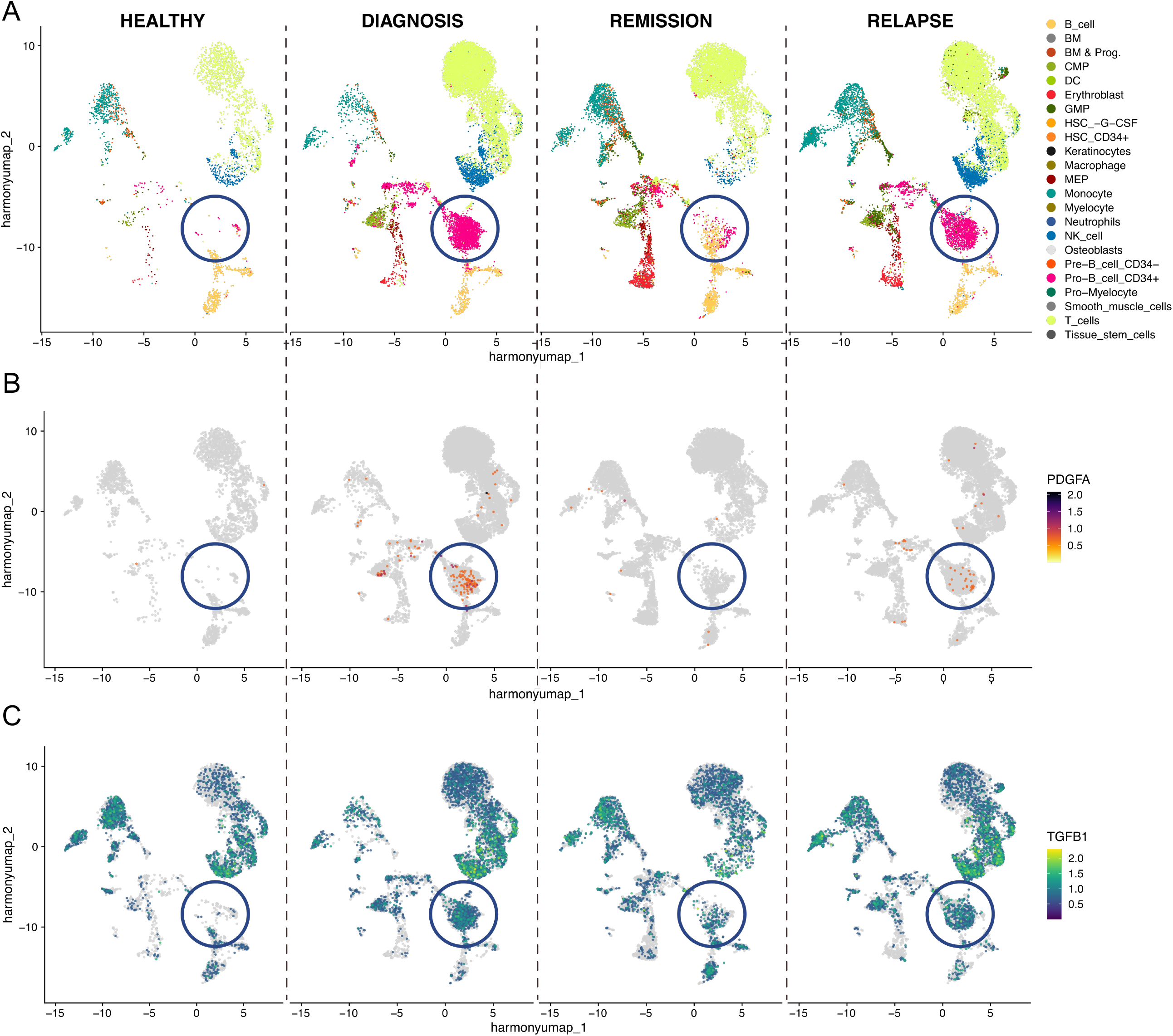
Re-analysis of the scRNAseq dataset by Witkowski et al. [18]. UMAPs showing primary BM samples from healthy controls (n=4, left panel)) and B-ALL patients (ETV6::RUNX1, n=5; Ph+, n=2) at different stages of the disease. **(A)** cell clusters were annotated based on Human Primary Cell Atlas Data. Clustering annotation was done via SingleR. Cell types are indicated by different colors (legend). **(B, C)** Expression of PDGFA **(B)** and TGFB1 transcript levels **(C).** Relative expression levels are indicated in the color legend. Localization of the main ALL blast cell population is indicated by the blue circle.

### Spatial expression of TGFB1 and PDGFA RNA

Since we observed an increased expression of *TGFB1* and especially *PDGFA1* in ALL samples compared to PMF and controls we wanted to investigate the spatial expression of these mRNAs further. Images were processed in Aviris to create T*GFB1* and *PDGFA1* objects, which illustrated high mRNA expression especially in ALL MKs (Fig 5.). We then calculated the surface ratio of *TGFB1* and *PDGFA1* expressing objects in small (2,86 mm^2^) representative MK regions to same-sized regions not containing MKs (Fig. 5 A-E). Here, the data showed that *PDGFA1* expression was relatively similar in MK regions compared to non-MK regions in ALL samples with fibrosis grade of 0 or 1, PMF, and control samples. In contrast, however, ratios were higher in ALL samples with fibrosis grade 2 (Fig. 5 F), indicating increased *PDGFA1* expression in fibrotic ALL MKs. A similar pattern was recorded for *TGFB1* ratios, i.e. they were higher in ALL with grade 2 fibrosis compared to ALL fibrosis 0, PMF and controls (Fig. 5 G).

**Figure 5.**
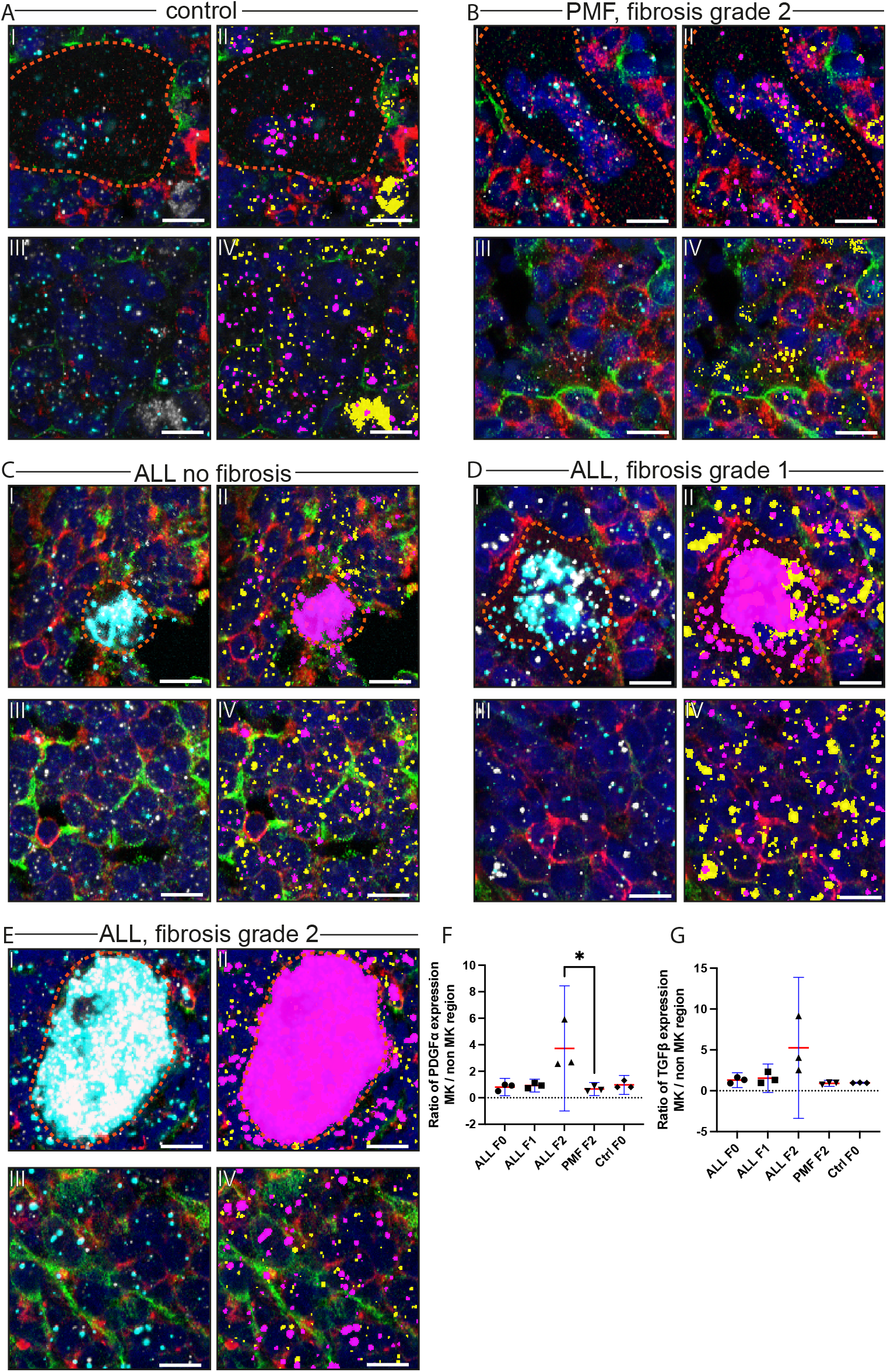
TGFB1 and PDGFA1 spatial expression analysis in ALL, PMF, and control samples. (A-E): Five-channel overlay of RNAscope staining together with IF antibody staining shown as object generated in Arivis. Nuclei were stained with DAPI (blue), CD271 with secondary-AF488 in green staining MSCs, CD45 with secondary-AF700 in red staining for hematopoietic cells, RNAscope® probe for TGFB1-C2 coupled to TSA Vivid™ 650 in cyan, and RNAscope® probe for PDGFA1-C1 coupled to TSA Vivid™ 570 in gray. (A): control sample C1 (B): PMF sample F2 (fibrosis grade 2) (C): ALL sample P1 (fibrosis grade 0) (D): ALL sample P5 (fibrosis grade 1) (E): ALL sample P7 (fibrosis grade 2). (I-II): MK areas are marked by the orange dotted line, (III-IV): areas without MKs. (II, IV): overlayed objects for TGFB1 (Pink) and PDGFA1 (yellow) generated with Arivis. Scale bars indicate 10 µm. (F-G): Ratios of mean cytokine mRNA positive surface areas [μm^2^] in MK to non-MK regions. (F) shows data for PDGFA1, (G) shows data for TGFB1. Individual data points are shown as symbols. Means are indicated as red bars with standard deviation (blue). Statistical analysis was performed using the Kruskal-Wallis multiple comparisons test (*: p ≤ 0.05).

Analysis of *TGFB1* and *PDGFA1* expression in individual cells confirmed these finding (Figure 6, supplementary Table 1). MKs showed high cytokine expression levels in all samples. *PDGFA1* expression in CD45^+^ cells was similar to CD45^-^CD271^-^ cells in ALL samples, whereas it was lower in the control and PMF (Figure 6 A). *PDGFA1* levels in CD271^+^ cells were lower compared to other cell types in all samples, except for ALL with fibrosis grade 2. Similar expression patterns were observed for *TGFB1* (Figure 6B). When comparing the different cell types, highest expression levels of *TGFB1* and *PDGFA1* were observed in high-fibrosis ALL-MKs (suppl. Table 1). *PDGFA1* levels in PMF MKs were also higher compared with controls and non-fibrotic ALL. Furthermore, cytokine expression levels were higher in CD271^+^ cells from ALL patients with fibrosis compared with PMF and controls. Expression levels in other cell types did not show a clear pattern (suppl. Table 1).

**Figure 6.**
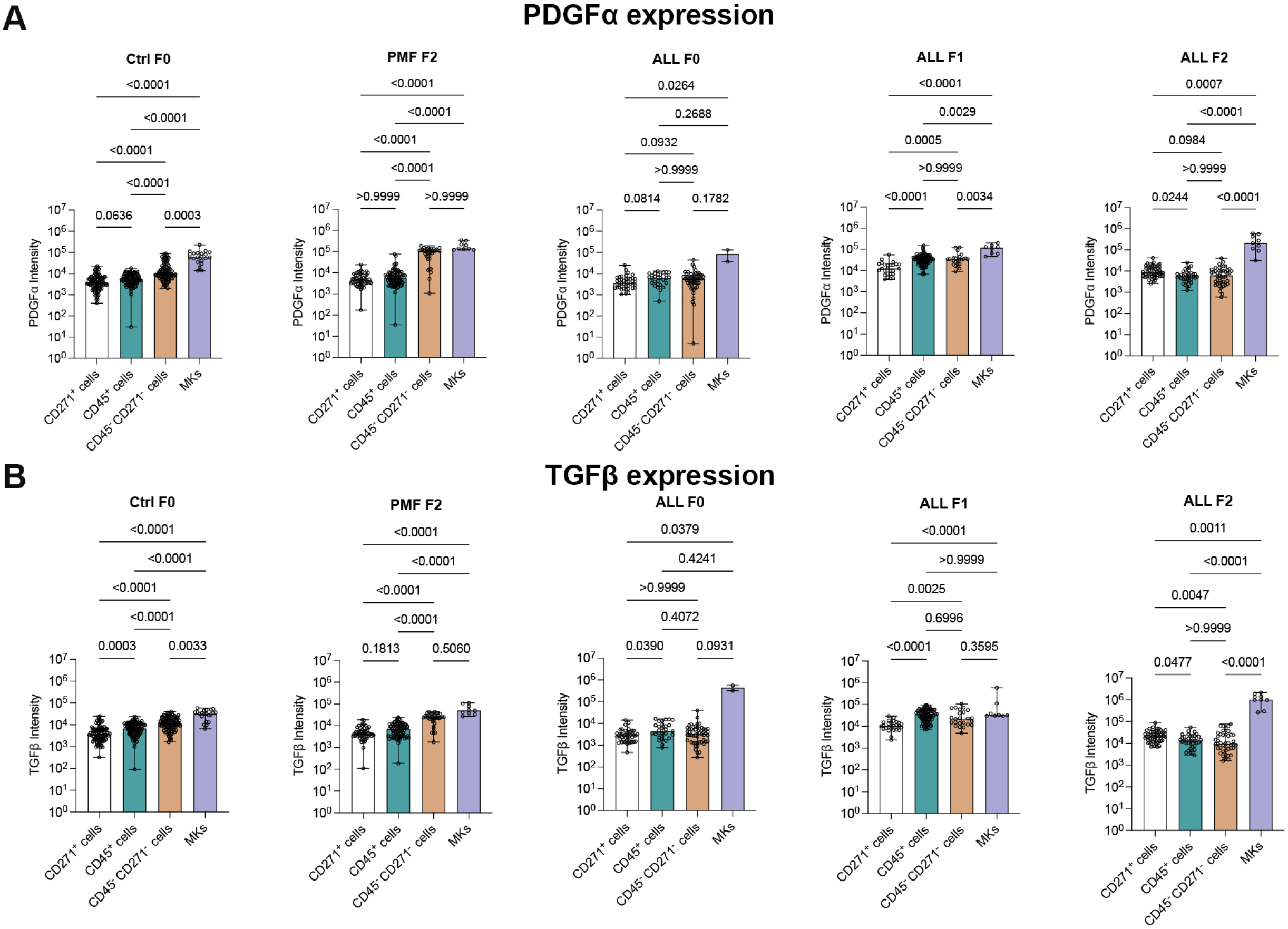
Quantitative in situ expression analysis of TGFβ and PDGFα. PDGFα and TGFβ intensities of single CD271^+^ cells (A), CD45^+^ cells (B), CD45^-^CD271^-^ cells (C), and megakaryocytes (MKs) (D) were measured on control sample C1 (Ctrl F0, no fibrosis), PMF with fibrosis grade 2 (PMF F2, sample ID F1), ALL with fibrosis grade 0 (ALL F0, sample ID P1), ALL with fibrosis grade 1 (ALL F1, sample ID P5), and ALL with fibrosis grade 2 (ALL F2, sample ID P7). Data are given as indiviual measurements in single cells (open circles), median of the sum intensity of individual cells (bars), and range (error bars). Statistical analysis was performed using the Kruskal-Wallis test by comparing pairs of cell types as indicated by the horizontal lines. P-values are provided above the lines.

## Discussion

Recent developments in tissue analysis technologies have opened new avenues to gather important information on complex cellular composition and gene/protein expression in a spatial matter. Modern spatial omics technologies thus have the potential to provide important novel insights into disease mechanisms by directly putting disease-related changes in their relevant cellular context In this study, we developed a novel and advanced analysis method by combining multi-color immunofluorescence staining with in-situ RNA expression analysis (RNAscope^®^) to investigate differences in cellular composition and spatial expression of two important fibrotic drivers, *TGFB1* and *PDGFA1*, in the BM of patients with acute lymphoblastic leukemia, ALL.

Our study is motivated by the clinical finding that fibrosis in pediatric ALL has been linked to unfavorable outcomes as reported by Norén-Nyström et al. [3]. These authors evaluated fiber reticulin density in 166 diagnostic bone marrow samples from a cohort of a total of 433 children with ALL. To our knowledge, this is the largest study addressing this question and the data demonstrated that reticulin fiber density (RFD) at diagnosis correlated to the levels of minimal residual disease (MRD) at day 29 of treatment. Patients with high MRD levels had higher RFD and B-ALL patients with low RFD at diagnosis and a rapid reduction of RFD on day 29 had a favorable outcome compared to patients with the same baseline RFD level at diagnosis but a slow RFD reduction.

It is important to mention that biopsy frequency was high in this pediatric cohort of patients, which is due to the fact that bone marrow biopsies in children are usually taken under anesthesia and that pediatric ALLs in Sweden are treated within the same study protocol. However, routines might differ between countries and the situation is completely different in adults where a biopsy is usually only taken in case of unsuccessful aspiration (“dry tap”).

We therefore chose to focus on pediatric B-ALL and combined our recently-established multi-color IF surface marker BM analysis [17] with in-situ expression analysis of *TGFB1* and *PDGFA1* mRNA expression using RNAscope^®^ technology. Because little is known about the mechanisms that drive fibrosis in acute leukemia, which likely is a secondary phenomenon, we chose to focus on fibrosis-related cytokines which are known key players in primary myelofibrosis, PMF.

Our advanced five-color panel immunofluorescence (IF) analysis method employs repetitive cycles of staining and bleaching, enabling the use of antibody combinations for surface marker detection that would otherwise be incompatible. Notably, the bleaching step not only eliminates the fluorochrome signal but also likely induces structural changes in the antibody, preventing secondary antibody binding. This allows subsequent staining with another primary antibody from the same species and same isotype [17].

The five-color panel bone marrow scanning of standard pathology FFPE samples from ALL, PMF, and controls recapitulated known pathological features. PMF biopsies showed hypercellularity and increased MK numbers forming dense clusters with atypical morphology. The PMF samples thus showed previously described atypical MK features [30, 31] and also high levels of CD271 expression with extensive fibrillary networks and increased vessel density, both of which are characteristics of myelofibrosis [32]. ALL samples on the other hand showed the typical picture of high-frequency uniform blast cells and a reduced number of megakaryocytes [7, 9].

We then explored the density of mesenchymal stromal cells (MSCs) identified by their CD271 expression. MSCs play a pivotal role in the HME and can contribute to fibrosis by cytokine-induced proliferation and differentiation into myofibroblasts [33]. We observed an increase in CD271^+^ MSCs in both fibrotic ALL and PMF samples. Notably, we observed a dense CD271 expression pattern close to blood vessels, mirroring the distribution of fibrotic regions as seen in conventional staining. Increased CD271^+^ MSC volumes, particularly in ALL samples with higher fibrosis degrees, imply a potential association between MSC density and fibrosis progression. Of note, there is evidence from published data that leukemic cells can produce stromal growth factors, like basic fibroblast growth factor (bFGF) and PDGFs, that can lead to fibrosis progression [34, 35]. However, a spatial analysis of fibrosis-related cytokine production in ALL marrows has not been reported thus far.

In the next step, we therefore examined the spatial expression of the two key fibrotic drivers *TGFB1* and *PDGFA1* which are implicated in fibrotic processes, including the activation of fibroblasts and the promotion of extracellular matrix deposition [36] [37]. In our study, we found increased expression of *TGFB1* and *PDGFA1* in leukemic blasts and especially high expression in MKs in fibrotic ALL BMs in comparison with PMF, which was confirmed by analyzing expression in single cells. This finding is novel and surprising as MKs in ALL – in contrast to PMF – are not part of the malignant clone and it clearly suggests a potential role of these fibrotic drivers in the fibrotic process in the context of ALL.

Expression of *TGFB1* and *PDGFA1* in ALL blasts was also confirmed by reanalysis of the single cell data set published by Witkowski et al. [18]. Here, our analysis showed that *TGFB1/PDGFA1* double-positive blasts were clearly detectable at diagnosis, they were absent in remission samples and reemerged in relapse samples. Although this data set showed interesting results in the B-ALL blast compartment it did not allow an investigation of MKs and CD45 negative cells as these were excluded during sample preparation. Also, information on fibrosis grade was not provided in this publication. Nevertheless, the dataset was identified by us as best suited of the currently available reports to address our questions. Naturally, a better tailored single cell approach also considering fibrosis grade and including all fibrosis-relevant cell populations would be certainly important to pursue in future experiments.

We found an elevated expression of fibrosis-relevant factors in ALL and especially a high expression of *PDGFA1* in MKs of ALL patients with higher degrees of fibrosis, which points to a potential correlation between fibrosis severity and fibrotic pathway activity. This is the first time that potential fibrosis mechanisms in ALL have been identified on the single cell level and, certainly, is an intriguing observation which will be followed up functionally by using a novel BM fibrosis xenotransplantation model recently developed by us (unpublished data). Furthermore, our data clearly motivates further experimentation including a larger panel of fibrosis-relevant factors, e.g. by utilizing advanced methods for multiplexed in-situ quantification of relevant BMSC protein and RNA targets, such as the Xenium platform (10x Genomics) or extended RNAscope multiplexing approaches. Last, longitudinal studies correlating treatment response and expression of fibrotic drivers and BM fibrosis levels could provide further and important information regarding the potential clinical significance and potential therapeutic options to reverse fibrosis.

Understanding the involvement of fibrotic drivers like *TGFB1* and *PDGFA1* in the disease process could offer new avenues for future therapeutic interventions. Targeting these pathways may provide opportunities for developing anti-fibrotic strategies to improve treatment response and, thus, the overall prognosis for patients with ALL. Early intervention to inhibit or reverse the fibrotic process might potentially overcome the negative effects of fibrosis on treatment outcomes. Moreover, investigating the crosstalk between leukemic cells, stromal cells, and fibrotic drivers could uncover novel targets for combination therapies that address both the hematological and microenvironmental components of the disease.

In summary, by investigating the spatial expression of the fibrotic drivers *TGFB1* and *PDGFA1*, as well as the density of CD271^+^ MSCs, we uncovered – for the first time – potential associations between fibrosis, microenvironmental components, and abnormal cytokine production in ALL. Our data indicate that the same fibrosis-driving mechanisms are operative in two otherwise unrelated diseases, i.e. ALL and PMF, thus providing important novel insights into the potential pathomechanisms that drive fibrosis in acute lymphoblastic leukemias.

## Ethics Approval

The sampling of patient material and sample preparation was approved by the institutional review committee in Lund (Regionala Etikprövningsnämnden i Lund, approvals no. 2014/776 and 2018/86) and all procedures were performed following the Helsinki Declaration of 1975, as revised in 2008.

## Author Contribution Statement

S.B. designed and performed experiments, analyzed data, and wrote the manuscript. C.D. performed experiments and analyzed data. D.N.V performed the scRNA seq data reanalysis and contributed to writing the manuscript. R.F. and A.L.S performed experiments, analyzed data, and contributed to wroting the manuscript. A.H. provided critical patient samples and contributed to writing the manuscript. J.E. provided critical patient samples, contributed to data analysis, and performed pathological evaluations. H.L. co-designed experiments and contributed to writing the manuscript. S.S. supervised the study, designed the experiments, and wrote the manuscript. All authors read and approved the final paper.

## Data Availability Statement

n/a

## Acknowledgment

The authors would like to thank the Lund Stem Cell Center Imaging facility personnel for technical assistance. The Lund University Bioimaging Centre (LBIC), Lund University, is gratefully acknowledged for providing experimental resources. We gratefully acknowledge Cristina-Daria Ciornei Karlsson for supplying us with high-quality BM biopsy cuts.

