## Supplementary material for "Combined multi-color immunofluorescence staining and spatial in situ mRNA expression analysis identifies potential fibrosis drivers in acute lymphoblastic leukemia": Table 1

**Table 1: Patient characteristics.**

| **ALL patients** |  |  |  |  |  |
| --- | --- | --- | --- | --- | --- |
| **Sample ID** | **Diagnosis** | **(WHO 2022 subtype)** | **Fibrosis grade** | **Sex** | **Age** |
| **P1** | B-ALL | with other defined genetic alterations. Fusion IGH/DUX4 | 0 | F | 2 |
| **P2** | B-ALL | with ETV6::RUNX1 fusion | 0 | M | 5 |
| **P3** | B-ALL | with other defined genetic alterations. TCF3: ZNF384 | 0 | F | 7 |
| **P4** | B-ALL | NOS | 1 | M | 3 |
| **P5** | B-ALL | with other defined genetic alterations. Fusion IGH/DUX4 | 1 | M | 5 |
| **P6** | B-ALL | High hyperdiploidity | 1 | M | 4 |
| **P7** | B-ALL | ETV6::RUNX1 fusion | 2 | M | 4 |
| **P8** | B-ALL | Unknown | 2 | M | 3 |
| **P9** | B-ALL | High hyperdiploidity | 1 to 2 | F | 3 |
| **PMF patients and normal controls** | | | | | |
| **Sample ID** | **Diagnosis** | | **Fibrosis grade** | **Sex** | **Age** |
| **F1** | PMF |  | 2 | F | 75 |
| **F2** | PMF |  | 2 | F | 32 |
| **F3** | PMF |  | 2 | M | 47 |
| **C1** | Normal control | | 0 | M | 50 |
| **C2** | Normal control | | 0 | F | 80 |
| **C3** | Normal control | | 0 | F | 76 |
| Abbreviations: ALL, acute lymphoblastic leukemia; PMF, primary myelofibrosis; M, male; F, female; NOS, not otherwise specified | | | | | |
