## Supplementary materials for "Combined multi-color immunofluorescence staining and spatial in situ mRNA expression analysis identifies potential fibrosis drivers in acute lymphoblastic leukemia"

**Bräunig et al.: Supplementary Figure 1**

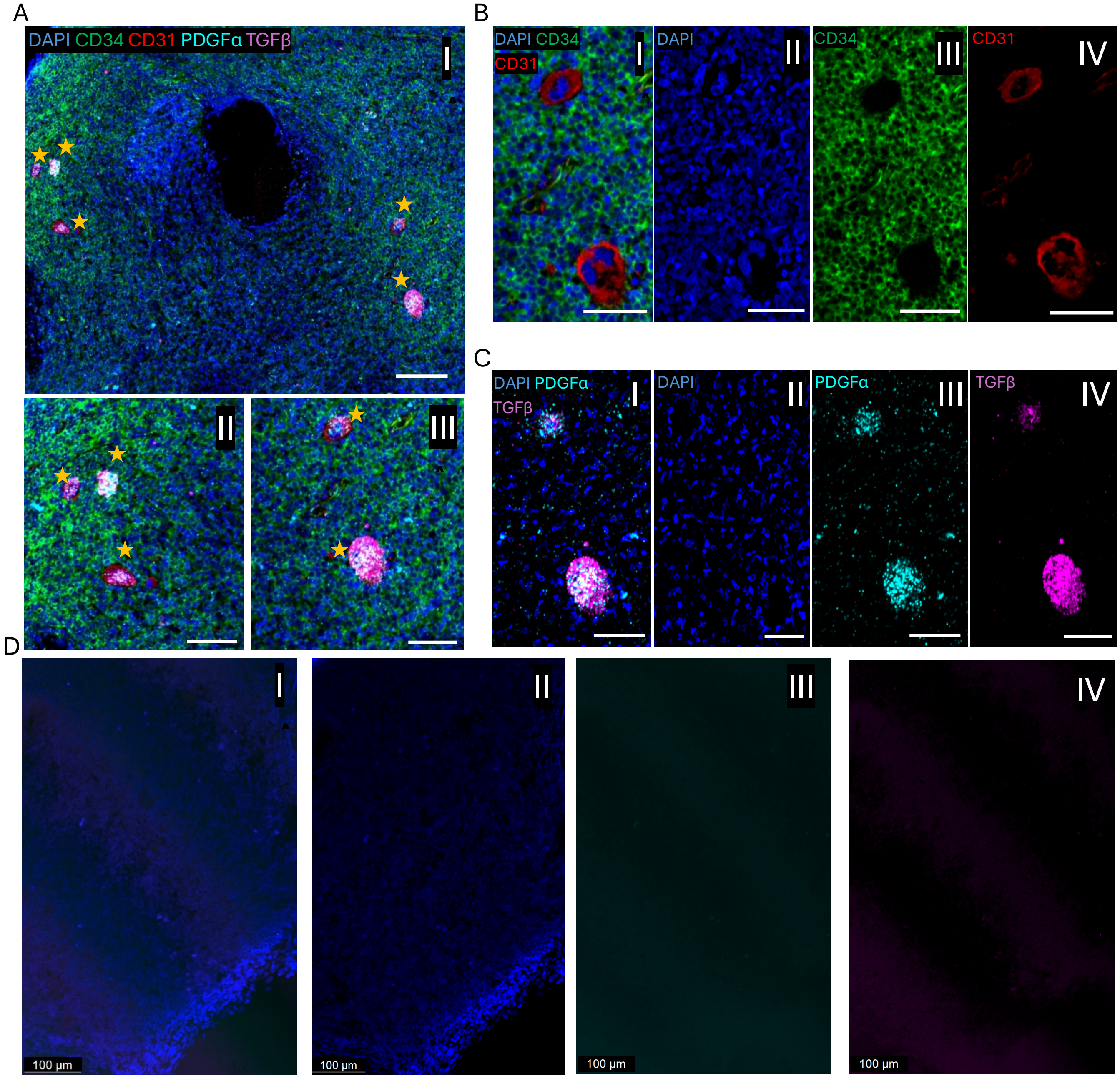


***Supplementary Figure 1: Megakaryocyte identification and in situ expression of TGFB1 and PDGFA1 in ALL bone marrows****. (A) Overview (I, scale bar indicates 100 µm) and zoom-in (II and III, scale bars indicate 50 µm) overlay of RNAscope and IF antibody stainings in consecutive slides of ALL sample P7 (fibrosis grade 2). Megakaryocytes (MKs) are indicated by yellow stars. (B) Three-channel overlay of IF antibody staining (I); nuclei were stained with DAPI (II); primary anti-CD34 antibody stained with secondary antibody (AF-488, green) (III); and primary anti-CD31 antibody stained with secondary antibody (AF-568, red), for ECs and MKs (IV). Scale bars indicate 50 µm. (C) Three-channel overlay of TFGB1 and PDGFA1 mRNA in situ hybridization (I); nuclei were stained with DAPI (II); RNAscope® probe for PDGFA1-C1 coupled to TSA Vivid™ 570 in cyan (III); and RNAscope® probe for TGFB1-C2*

*coupled to TSA Vivid ™ 650 in magenta (IV). Scale bars indicate 50 µm. (D) Three channel overlay of RNAscope® staining with the negative control probes targeting the DapB gene (accession # EF191515) of the Bacillus subtilis strain SMY. Nuclei were stained with DAPI (blue). Negative control with DAPI alone (II); negative control probe plus TSA Vivid™ 570 in cyan (III) and TSA Vivid ™ 650 in magenta (IV), respectively. Scale bars indicate 100 µm.*

**Bräunig et al.: Supplementary Figure 2**

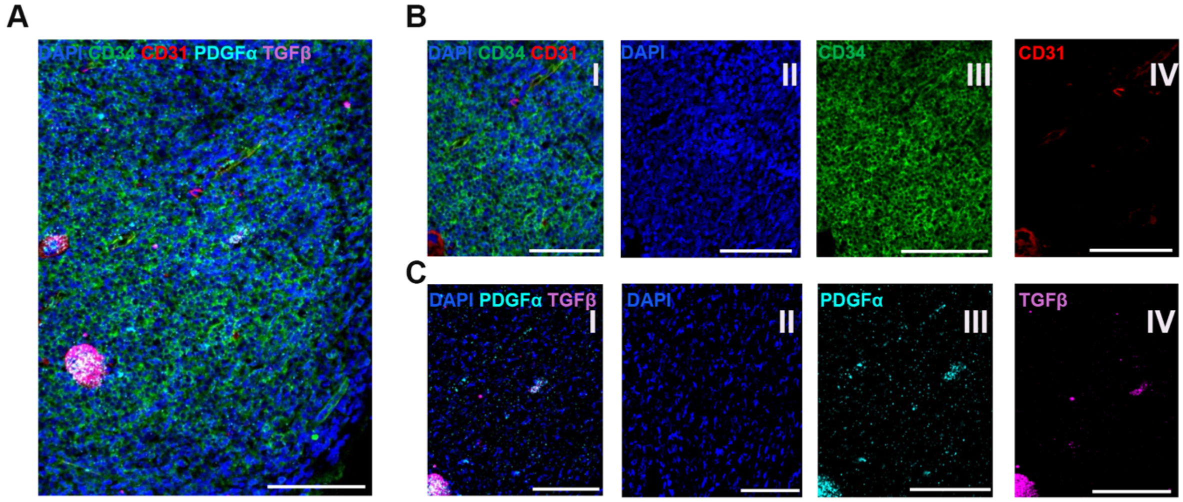


***Supplementary Figure 2: Blast identification and in situ expression of TGFB1 and PDGFA1 in ALL bone marrows.*** (A) Overview of RNAscope and IF antibody staining overlays from consecutive bone marrow slides of an ALL patient with fibrosis grade 2 (sample P7). (B) Three-channel overlay of IF antibody plus DAPI staining (I); nuclei were stained with DAPI (II); CD34 with secondary AF-488, in green, identified ALL blasts (III); and CD31 with secondary-AF568, in red, for identification of ECs and MKs (IV). (C) Three-channel overlay of TGFβ and PDGFα mRNA in situ hibridization (I); nuclei were stained with DAPI (II); RNAscope® probe for PDGFA1-C1 coupled to TSA Vivid™ 570 in cyan (III); and RNAscope® probe for TGFB1-C2 coupled to TSA Vivid ™ 650 in magenta (IV). Scale bars indicate 100 µm.

***Supplementary Table 1:***

**A**

**CD271$^+$ cells**

| | Ctrl F0 | PMF F2 | ALL F0 | ALL F1 | ALL F2 |
|---|---|---|---|---|---|
| Counted cell numbers | 100 | 53 | 43 | 26 | 53 |
| PDGFα | 3865 | 4308 | 3496 | 12370 | 8427 |
| TGFβ | 3631 | 4111 | 2806 | 10359 | 20268 |

**B**

**CD45$^+$ cells**

| | Ctrl F0 | PMF F2 | ALL F0 | ALL F1 | ALL F2 |
|---|---|---|---|---|---|
| Counted cell numbers | 100 | 84 | 29 | 83 | 36 |
| PDGFα | 5412 | 5884 | 6141 | 34157 | 5740 |
| TGFβ | 6800 | 7034 | 4324 | 35776 | 13458 |

**C**

**CD271$^-$ CD45$^-$ cells**

| | Ctrl F0 | PMF F2 | ALL F0 | ALL F1 | ALL F2 |
|---|---|---|---|---|---|
| Counted cell numbers | 100 | 36 | 54 | 24 | 43 |
| PDGFα | 10062 | 102912 | 5575 | 33182 | 6228 |
| TGFβ | 11732 | 24061 | 3413 | 22260 | 9401 |

**D**

**MK cells**

| | Ctrl F0 | PMF F2 | ALL F0 | ALL F1 | ALL F2 |
|---|---|---|---|---|---|
| Counted cell numbers | **21** | **9** | **2** | **8** | **2** |
| PDGFα | 59718 | 140544 | 82801 | 118150 | 210592 |
| TGFβ | 31541 | 50065 | 439038 | 33041 | 1001100 |

**Supplementary Table 1.** PDGFα and TGFβ intensity of single CD271$^+$ cells (A), CD45$^+$ cells (B), CD45$^-$CD271$^-$ cells (C), and megakaryocytes (D) were measured on control sample C1 (Ctrl F0, no fibrosis), PMF with fibrosis grade 2 (PMF F2, sample ID F1), ALL with fibrosis grade 0 (ALL F0, sample ID P1), ALL with fibrosis grade 1 (ALL F1, sample ID P5), ALL with fibrosis grade 2 (ALL F2, sample ID P7). Data are given as the median of the sum intensity of individual cells. For patient characteristics and sample ID, please see Table 1.
